# The Metabolomics Workbench File Status Website: A Metadata Repository Promoting FAIR Principles of Metabolomics Data

**DOI:** 10.1101/2022.03.04.483070

**Authors:** Christian D. Powell, Hunter N.B. Moseley

## Abstract

**Motivation:** An updated version of the mwtab Python package for programmatic access to the Metabolomics Workbench (MetabolomicsWB) data repository was released at the beginning of 2021. Along with updating the package to match the changes to MetabolomicsWB’s ‘mwTab’ file format and to enhance the package’s functionality, the package included format validation facilities which were used to detect and catalog file inconsistencies and errors across all publically available datasets in MetabolomicsWB.

**Results:** The Metabolomics Workbench File Status website was developed to provide continuous validation of MetabolomicsWB data files and a useful interface to find all inconsistencies and errors that are found. This list of detectable issues/errors include format parsing errors, format compliance issues, access problems via MetabolomicsWB’s REST interface, and other small inconsistencies that can hinder reusability. The website uses the mwtab Python package to pull down and validate each available analysis file and then generates an html report. The website is updated on a weekly basis. Moreover, the Python website design utilizes GitHub and GitHub.io, providing an easy to replicate template for implementing other metadata, virtual, and meta-repositories.

**Availability:** Metabolomics Workbench file status website can be accessed at: https://moseleybioinformaticslab.github.io/mwFileStatusWebsite/. The mwtab Python library is available on GitHub: https://github.com/MoseleyBioinformaticsLab/mwtab, PyPI: https://pypi.org/project/mwtab/, and documentation is available on ReadTheDocs: https://mwtab.readthedocs.io/. Metabolomics Workbench analysis data files used for analysis presented here along with the generated HTML files of the website are available on FigShare: https://doi.org/10.6084/m9.figshare.19221159.

**Supplementary information:** Supplementary data are available at *FigShare* online.

## 1 Introduction

In 2006, the Congress of the United States of America passed the National Institutes of Health (NIH) Reform Act of 2006 to reauthorize and reorganize the NIH, which also established the NIH Common Fund to support cross-cutting, trans-NIH programs which are involved in two or more of the NIH’s Institutes or Centers (ICs). Today, the Common Fund consists of more than 30 active projects and has funded more than 50 projects in total (National Institute of Health, 2022). One currently funded project is the Metabolomics project to inform basic, translational, and clinical research. The Common Fund’s Metabolomics project established the Metabolomics Workbench (MetabolomicsWB) as a longstanding, public repository for national and international metabolomics data and metadata (Sud *et al.,* 2016). Today, MetabolomicsWB is host to more than 1,600 studies with over 2,500 individual metabolomics analyses currently available.

In 2018, the mwtab Python package was initially released with the intent of providing a ‘pythonic’ means of access and manipulation of data and metadata hosted on MetabolomicsWB (Smelter and Moseley, 2018). The package was developed with the intention of promoting the FAIR principles of data to be findable, accessible, interoperable, and reusable (Wilkinson *et al.*, 2016; Boeckhout *et al.,* 2018). Along with the release of the mwtab Python package, the package was used to find a number of inconsistencies between the mwTab file format specification and actual mwTab data files hosted by MetabolomicsWB.

With the release of the 1.1.0 version of the mwtab Python package (Powell and Moseley, 2021), the MetabolomicsWB file validation functionality of the package was greatly improved and updated to follow the latest mwTab file format specification. The mwtab package was again used to find formatting errors in mwTab files, undercovering that a majority of the datafiles hosted on MetabolomicsWB have some degree of inconsistency to their file format specification.

In order to make individual researchers wishing to use data from MetabolomicsWB aware of possible issues with specific datasets, the MetabolomicsWB File Status website was developed. The website uses the mwtab Python package to provide a simple way to determine which analysis files in the MetabolomicsWB contain file formatting errors, inconsistencies, or are simply unavailable through MetabolomicsWB’s REST interface.

## 2 Methods

The Metabolomics Workbench (MetabolomicsWB) file status website is available through GitHub Pages. All scripts and files used to generate the website are made open-source and are publicly available within the GitHub repository hosting the website.

### 2.1 The Metabolomics Workbench File Status Website

The MetabolomicsWB file status website is a series of HTML pages hosted via GitHub Pages. The HTML files are regularly updated using a series of Python scripts hosted in the same GitHub repository with the GitHub Pages deployment. The scripts extensively use the mwtab Python package to collect and validate the available analysis data files from MetabolomicsWB. The first script to be run is the ~mwFileStatusWebsite.validator.py script. The script uses the ~mwtab.mwrest._pull_study_analysis() method to generate a dictionary containing a mapping of all available study IDs and their corresponding analysis IDs. The ~mwtab.read_files() method is then used iteratively to retrieve each available analysis data file in mwTab (plain text) and JSON format. The ~mwtab.validate_file() method is then used to generate a validation log file and collect the validation status of each analysis file. Along with validating each individual data file, the mwTab and JSON file formats are compared to ensure consistency between the two. The ~mwFileStatusWebsite.compare.py script contains a series of methods for performing these comparisons based on an updated version of the comparison analysis previously performed (Powell and Moseley, 2020). The final script to run is the ~mwFileStatusWebsite.generate_html.py script which contains methods for taking the collected validation and comparison statistics and inserting them into HTML templates to form the MetabolomicsWB file status website. The final templates, along with a CSS style, is then pushed to the GitHub repository and the webpage is automatically updated.

### 2.2 The mwtab Python Package Version 1.2.4

The mwtab Python3 package was updated alongside the MetabolomicsWB file status website. The most significant changes to the package include the restructuring of validation logs generated when using the ~mwtab.validator.validate_file() method. The method now has the option to return a structured string validation log. This change was included to allow for the generation of discrete validation logs for each available analysis data file in both mwTab (plain text) and JSON formats. Upon validation, the validation log string is captured and saved into a plaintext file which is then uploaded to the GitHub repository hosting the MetabolomicsWB file status website.

## 3 Results

The Metabolomics Workbench (MetabolomicsWB) file status validation website is available on GitHub Pages at: https://moseleybioinformaticslab.github.io/mwFileStatusWebsite/. The source code for the Python scripts used to generate the webpage and validation logs for each available mwTab data file are available in the same GitHub repository that is used to host the webpage: https://github.com/MoseleyBioinformaticsLab/mwFileStatusWebsite. The mwtab Python library which is used to validate the analyses from MetabolomicsWB is available on GitHub: https://github.com/MoseleyBioinformaticsLab/mwtab, PyPI: https://pypi.org/project/mwtab/, and documentation is available on ReadTheDocs: https://mwtab.readthedocs.io/. The analysis data files used for this analysis were downloaded on February 19th, 2022 and the generated HTML files of the website are available for download at: https://doi.org/10.6084/m9.figshare.19221159.

### 3.1 File Errors in Metabolomics Workbench Analysis Datafiles

The ~mwtab.mwtab.validator.py module from the 1.2.4 version of the mwtab Python package was used to determine errors in both the ‘mwTab’ and JSON file formats for each available analysis data file. For the ‘mwTab’ formatted analyses, 1 file was inaccessible through MetabolomicsWB’s REST API, 303 files contained gross formatting errors preventing them from being parsed, 626 files contained minor formatting errors which were inconsistent with MetabolomicsWB’s ‘mwTab’ file format specification, and the remaining 1822 ‘mwTab’ formatted files passed all validation. For the JSON formatted analyses, 1 file was inaccessible through MetabolomicsWB’s REST API, 56 files contained gross formatting errors preventing them from being parsed, 951 files contained minor formatting errors which were inconsistent with MetabolomicsWB’s ‘mwTab’ file format specification, and the remaining 1744 JSON formatted files passed all validation.

### 3.2 File Consistency in Metabolomics Workbench Analysis Data Files

The ~mwFileStatusWebsite.mwFileStatusWebsite.compare.py module of the 1.0.0 version of the mwFileStatusWebsite Python3 package was used to compare the ‘mwTab’ (plain text with tab separation) and JSON formatted data files for each available analysis. Of the 2752 available analyses, 669 analyses had consistent ‘mwTab’ and JSON files, 1751 analyses had inconsistent ‘mwTab’ and JSON files, and 332 analyses could not be compared due to one or more of the file formats being unparsable or missing.

### 3.3 Development of Metadata, Virtual, and Meta-Repositories

First, a **metadata repository** is a repository type which provides additional data to enhance a pre-existing repository. For example, the MetabolomicsWB File Status website is technically a metadata repository that provides additional validation metadata to MetabolomicsWB analyses. Its implementation cleanly adds a GitHub-hosted Web User Interface (UI) to the validation metadata generated by the Python mwtab package. This easy-to-implement design can be duplicated for other metadata repositories as well as for virtual repositories. Second, a **virtual repository** is a repository type which only provides access to pre-existing data without adding novel data. An example of a virtual repository is Metabolome Exchange which provides access to datasets from four different metabolomics data repositories (Haug et al., 2017). Third, by combining both concepts and including extra data, this approach could be used to create a **meta-repository**, which overlays additional metadata onto a virtual repository that points to datasets, but also provides additional data access. Validated MetabolomicsWB datafiles are intentionally not provided by the MetabolomicsWB File Status website. Providing such files would shift the website from being a metadata repository to being a meta-repository. The MDACC Standardized Data Metabolomics Workbench Tool is a current example of a meta-repository which provides additional functionality allowing users to view study/analysis metadata from Metabolomics Workbench and download the metabolite data section separately (Casasent et al., 2022). Moreover, our approach would be ideal for efficiently implementing specialized public meta-repositories for specific scientific communities that utilize a well-established scientific repository for deposition, interoperability, and access functionality. Such meta-repositories are easier to implement and maintain than alternative full duplicate repositories like the PDB-REDO Database (Joosten et al., 2009), which contains re-analyzed versions of 3D macromolecular structure entries in the worldwide Protein Data Bank (Bermann et al., 2007).

**Fig 1.**
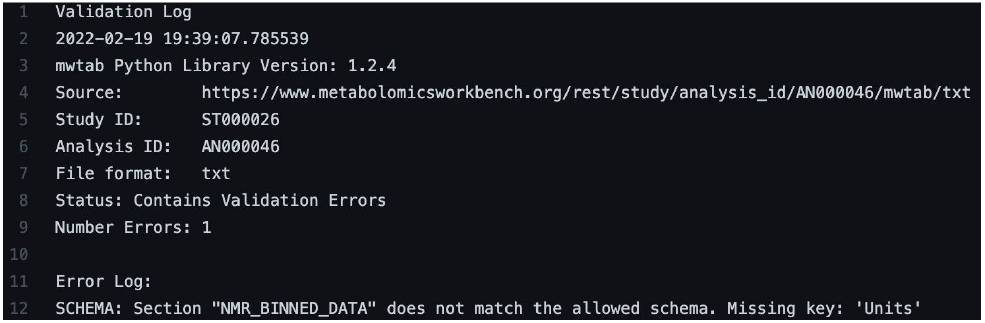
Example Validation Log Created by the mwtab Python Package. Validation logs are created by calling the ~mwtab.validate_file() method. The validation log consists of; metadata surrounding when and the conditions the validation was performed, metadata identifying the Metabolomics Workbench analysis data file the validation was performed on, and the number of validation errors collected if any.

**Fig. 2.**
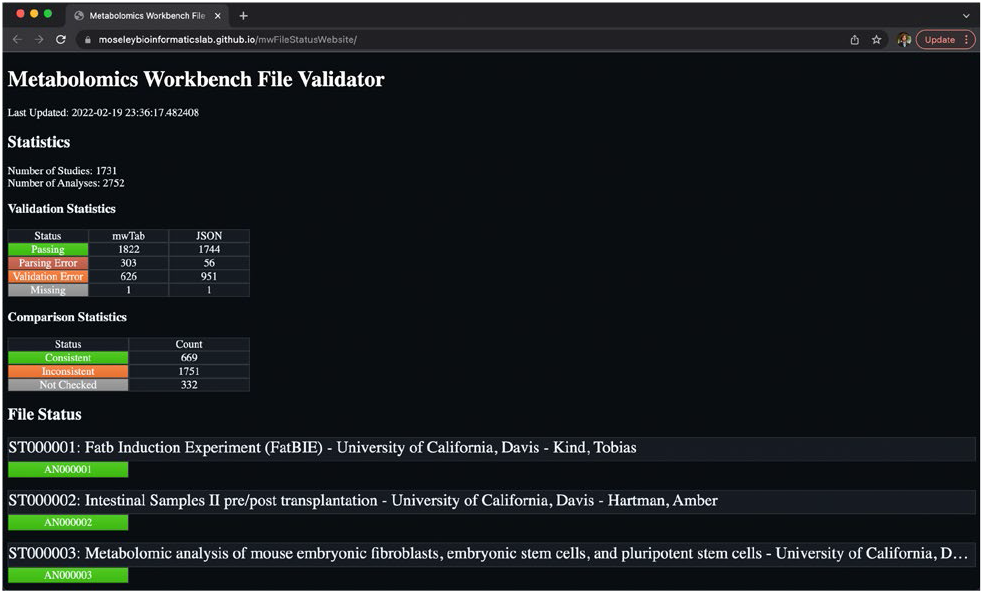
The Metabolomics Workbench File Status Website. The website uses Metabolomics Workbench’s dataset hierarchy of Studies and Analyses to logically display the file status of each file format of each analysis available from the Metabolomics Workbench data repository.

**Table 1.** File status for each file format of each analysis available from the Metabolomics Workbench data repository.^*a*^

| Status\File Format | mwTab | JSON |
| --- | --- | --- |
| Passing | 1822 | 1744 |
| Parsing Error | 303 | 56 |
| Validation Error | 626 | 951 |
| Missing | 1 | 1 |

<sup>a</sup> These results were generated on February 19th, 2022.

**Table 2.**
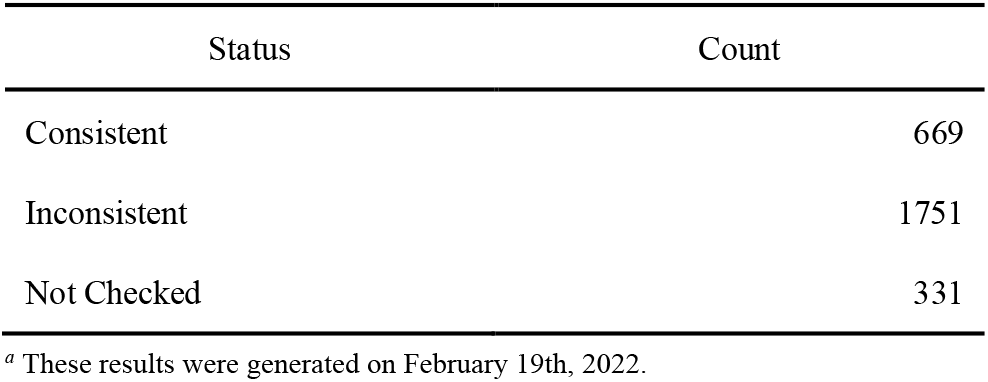
Comparison statistics for the comparison of the ‘mwtab’ (plain text) and JSON formatted data files of each analysis data file available from the Metabolomics Workbench data repository.^*a*^

## Acknowledgements

The authors would like to acknowledge the amazing degree of care and effort that Shankar Subramaniam, Eoin Fahy, and the whole MW/UC San Diego team have put into provisioning FAIR access to metabolite studies and their incredible effort in expanding and maintaining the repository. The authors would also like to acknowledge Nicholas Santini for his assistance in designing the HTML and CSS of the Metabolomics Workbench File Status website.

## Supplementary information

All the Metabolomics Workbench analysis data files which were available through MW’s REST interface have been uploaded to FigShare and are accessible at https://doi.org/10.6084/m9.figshare.19221159.

